# Profiling of Secondary Metabolites of Leaves of Indigenous and Introduced Grapes Collected from Hebron and Bethlehem Regions in the West Bank-Palestine

**DOI:** 10.64898/2026.02.09.704947

**Authors:** Jamil Harb, Thomas Hoffmann, Husameddin Isaid, Manal Shuaib, Abdullatif Husseini, Wilfried Schwab

## Abstract

Grape (*Vitis vinifera* L.) is one of the most cultivated plant species and has a long history in the Levant. Grape products, including leaves, are highly appreciated as healthy products, mainly because of their high levels of nutraceuticals. The significance of these products lies in the fact that poor diet is the primary cause of malnutrition, which is associated with severe noncommunicable diseases. Accordingly, this study aimed to profile secondary metabolites in a selection of grape genotypes from Hebron and Bethlehem regions in the West Bank-Palestine that include both indigenous and introduced genotypes. Fresh, delicate leaves from each genotype–region combination were analyzed for their content of secondary metabolites via LC–MS. The results revealed that the collection regions had a negligible impact, whereas the genotype impact was high and significant. More importantly, the secondary metabolites profiles of leaves allow for the clustering of the assessed genotypes into a few clusters, each with a specific set of metabolites that can serve as fingerprint profile. In conclusion, the results of the present study revealed the diversity of grape genotypes at the metabolomics level, which will help preserve indigenous grape genotypes and aid in the development of grape varieties that can cope with the adverse impacts of climate change.

## 1. Introduction

Grape (*Vitis vinifera* L.) is one of the most cultivated plant species worldwide, with a production volume of 72 million tonnes (FAO, 2024). With respect to the history of grapes, several reports (Dong et al., 2023) indicated the importance of the Western Asia domestication center in the evolution of grapevines approximately 11,000 years ago. In this sense, Goor (1966) claimed that the cultivation of European vines migrated from Anatolia to the Holy Land at approximately 5000 B.C. This long history led to significant variation in grape genotypes, with informal estimates suggesting the existence of more than 40 distinct genotypes in Palestine (PS).

Owing to the importance of grapevine leaves as a healthy foodstuff in most Mediterranean countries, research addressing their nutraceuticals is highly needed. In this respect, Kazemi and Ghasemi Bezdi (2022) reported that grape leaves could be used in lamb diets without deleterious effects. Furthermore, Maia et al. (2021) considered leaves a valuable addition to the diet. In a study conducted using grapevine leaves, Harb et al. (2013) reported that the leaves of the ‘Shami’ grape exhibited an anticancer effect. Furthermore, the enhancement of neurocognitive function in elderly individuals (Krikorian et al., 2012) and accelerated apoptosis in prostate cancer patients (Hudson et al., 2007) have been reported with various grape products. Another important aspect is the presumed tolerance of old indigenous grape genotypes to various biotic and/or abiotic stresses. This is highly relevant these days, with noticeable climate change (Seguin, 2010). In this respect, Palestinian farmers are concerned about shifts in seasonal rainfall patterns and further claim that a few old grape genotypes tolerate powdery mildew infection better. This reported tolerance of some strains to biotic and/or abiotic stresses might be the result of prolonged adaptation to harsh environments, particularly drought and high-temperature stresses. This tolerance of plant species to biotic and abiotic stresses has been reported to be closely associated with their metabolic profiles (Zandalinas et al., 2018). Apart from osmotic adjustment, a group of secondary metabolites, termed “stress-protective secondary metabolites,” might be involved in better adaptation to both biotic and abiotic stresses (Kaur and Ganjewala, 2019). Furthermore, the findings that plants that are tolerant to drought might also be more tolerant to UV radiation are valuable (Al-Oudat et al., 1998; Piri et al., 2011). This knowledge is highly valuable for cultivated plants that receive plenty of UV radiation in regions such as the Jordan Valley in PS (Robinson, 1848).

On the basis of the abovementioned scientific reports, this study aimed to elucidate the metabolic profiles of a set of major grape genotypes cultivated in PS. These findings will provide a basis for further studies that address potential correlations between metabolic profiles and the presumed tolerance to biotic and/or abiotic stresses. In addition, the data from metabolic profiling of grapevine leaves will be of significant benefit in promoting increased consumption of grapevine leaves as healthy foodstuffs, particularly in marginalized poor rural regions.

## 2. Materials and methods

### 2.1 Plant material

Fresh delicate, medium-sized grapevine leaves were collected in the spring season (May 2023) during the main flush period of vegetative growth. Fresh samples were collected from the following regions: Al-Khader (coordinates: 31°41′34″N 35°09′59″E (https://en.wikipedia.org/wiki/Al-Khader)), Halhul (coordinates: 31°34′44″N 35°05′57″E (https://en.wikipedia.org/wiki/Halhul)), and Hebron (coordinates: 31°31′43″N 35°05′49″E (https://en.wikipedia.org/wiki/Hebron))). All locations are in the southern part of the West Bank, PS. Leaves were collected from three separate vines as three biological replicates. The assessed genotypes are the following: Jandali, Marawi, Halawani, Baituni, Shami, Bairuti, and Eshuhki. Genotypes are identified on the basis of various parameters: during the collection period, farmers and local agricultural extension officers joined the collection team for the purpose of proper and accurate selection of genotypes. Extensive interviews were conducted with these farmers and officers about the collected genotypes, their characteristics, agronomic features, and tolerance/sensitivity to biotic and/or abiotic stresses. The necessity of this approach lies in the fact that no local scientific study that addresses grape genotypes and their morphological and agronomic characteristics and features is yet available. Directly after collection, the leaves were shock-frozen in liquid nitrogen and stored at −80°C until analysis.

The Institutional Review Board (IRB) approval was obtained from the research ethics committee at the Institute of Community and Public Health, Birzeit University before the beginning of the study (Supplementary file 1). Participation was voluntary, and the data collected were treated confidentially and anonymously.

### 2.2 Determination of antioxidant potential and polyphenol content

The total polyphenols were quantified via the Folin–Ciocalteu (FC) method: 0.125 g of finely ground leaf sample was dissolved in 70% ethanol, vortexed, incubated for 20 minutes, and further centrifuged for 15 minutes at 6000 rpm. The clear supernatants were centrifuged again at 10000 rpm. One hundred microliters of the clear supernatant was added to 200 µl of sodium carbonate solution (1 M), 500 µl of FC reagent (1 FC: 3 water (v/v)), and 200 µl of water. After 80 minutes of incubation in the dark, the absorbance was measured at 756 nm. A gallic acid solution (100 mg in 1 liter of 70% ethanol) was used for the calibration curve.

The antioxidant capacity was determined via the 2,2-diphenyl-1-picrylhydrazyl (DPPH) method. Samples were prepared by adding 0.25 g of finely ground powder (leaf samples) to 3 ml of 100% ethanol. The preparations were vortexed, sonicated for 20 minutes, and further centrifuged at 6000 rpm for 20 minutes. A total of 150 µl of the supernatant was added to 1000 µl of ethanol and centrifuged for 20 minutes at 10000 rpm. Finally, 500 µl of the clear supernatant was added to 500 µl of DPPH solution, which was subsequently vortexed and incubated for 30 minutes in darkness, after which the absorbance was ultimately measured at 515/517 nm. Trolox solution was used for the calibration curves.

### 2.3 Extraction and determination of individual secondary metabolites

Ten medium-sized mature leaves were collected from each grapevine, which represented one biological replicate. For each genotype/location combination, there were three biological replicates. Shock-frozen leaves were freeze-dried and further ground to powder. One hundred milligrams of the lyophilized tissue for each biological replicate was added to 500 μL of absolute methanol and 250 μL of the internal standard solution (50 mg biochanin A in 250 mL of absolute methanol). After vortexing for 1 minute and sonicating for 5 minutes, the mixed solutions were centrifuged at 13200 rpm for 10 minutes. The supernatants were collected in fresh tubes, and the pellets were extracted, vortexed, sonicated, and centrifuged once again as described above. The clear supernatants were pooled and placed in a SpeedVac for 2 hours. The dried residue from each replicate was dissolved in 35 μL of water, sonicated for 10 minutes, and finally centrifuged at 13200 rpm for 10 minutes. For each replicate, 20 μL of the clear supernatant was placed in an HPLC vial for final analysis via an LC–MS instrument (Agilent HPLC 1100; MS: Bruker Daltonics esquire3000 plus, Santa Clara, CA) that was equipped with a Phenomenex column (Luna 3u C18 (2) 100A′′, 150 mm × 2.0 mm (Part-Nr. 00F-4251-B0); Aschaffenburg, Germany). The instrumental analysis and data analysis of the chromatograms were the same as those reported by Harb et al. (2014).

### 2.4 Data analysis and statistics

The obtained data were subjected to statistical analysis using Costat software ((CoHort Software, Monterey, CA). FC and DPPH data were analyzed for significant differences at p values ≤ 0.05 via ANOVA and Tukey’s HSD test for mean separation. The equality of variances was assessed through Barttlet test using also Costat Software. Secondary metabolite profiling was achieved via R powered by our own algorithms. The statistical analysis for this set of data was conducted also using ANOVA test for significant differences at p values ≤ 0.05 (Supplementary data file 2).

## 3. Results

### 3.1 Antioxidant capacity and total (poly)phenol determination

Quantification of antioxidant capacity via the DPPH assay revealed pronounced and statistically significant differences (p < 0.001) among genotypes and locations (Table 1).

**Table 1:**
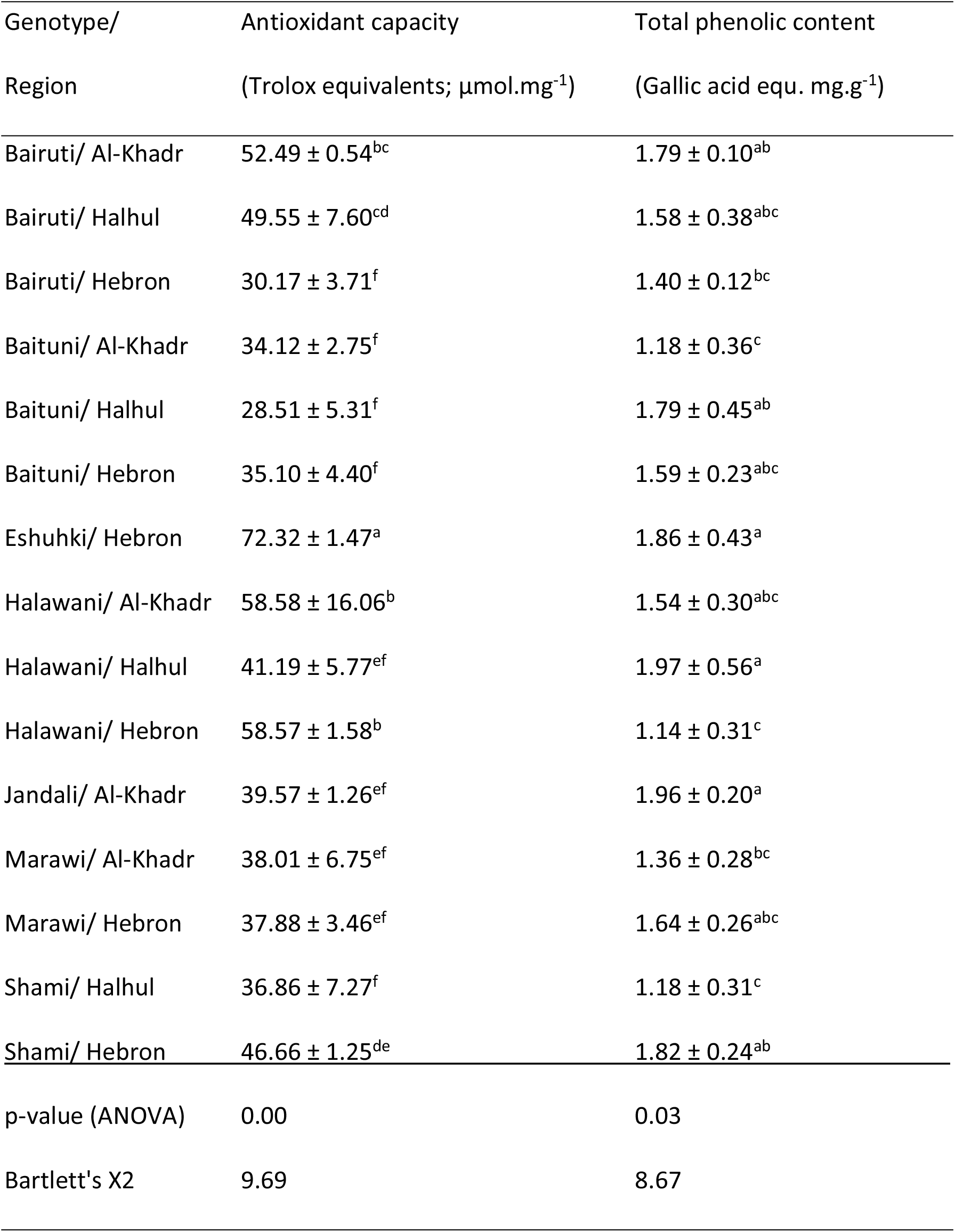
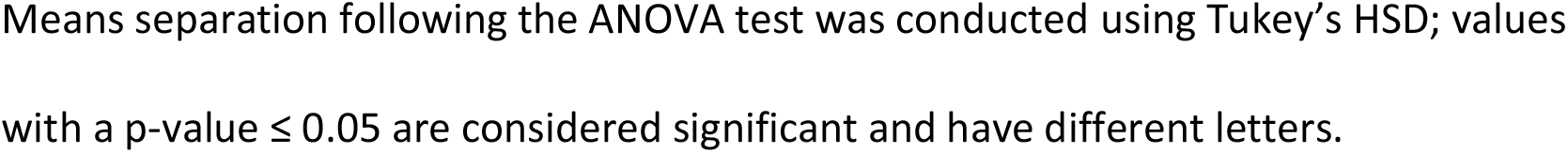
Antioxidant capacity (DPPH test) and total phenolic content (FC test) of the grapevine leaves from genotypes collected from various regions on the West Bank - PS.

| Genotype/<br>Region | Antioxidant capacity<br>(Trolox equivalents; $\mu\text{mol.mg}^{-1}$) | Total phenolic content<br>(Gallic acid equ. $\text{mg.g}^{-1}$) |
| --- | --- | --- |
| Bairuti/ Al-Khadr | $52.49 \pm 0.54^{bc}$ | $1.79 \pm 0.10^{ab}$ |
| Bairuti/ Halhul | $49.55 \pm 7.60^{cd}$ | $1.58 \pm 0.38^{abc}$ |
| Bairuti/ Hebron | $30.17 \pm 3.71^f$ | $1.40 \pm 0.12^{bc}$ |
| Baituni/ Al-Khadr | $34.12 \pm 2.75^f$ | $1.18 \pm 0.36^c$ |
| Baituni/ Halhul | $28.51 \pm 5.31^f$ | $1.79 \pm 0.45^{ab}$ |
| Baituni/ Hebron | $35.10 \pm 4.40^f$ | $1.59 \pm 0.23^{abc}$ |
| Eshuhki/ Hebron | $72.32 \pm 1.47^a$ | $1.86 \pm 0.43^a$ |
| Halawani/ Al-Khadr | $58.58 \pm 16.06^b$ | $1.54 \pm 0.30^{abc}$ |
| Halawani/ Halhul | $41.19 \pm 5.77^{ef}$ | $1.97 \pm 0.56^a$ |
| Halawani/ Hebron | $58.57 \pm 1.58^b$ | $1.14 \pm 0.31^c$ |
| Jandali/ Al-Khadr | $39.57 \pm 1.26^{ef}$ | $1.96 \pm 0.20^a$ |
| Marawi/ Al-Khadr | $38.01 \pm 6.75^{ef}$ | $1.36 \pm 0.28^{bc}$ |
| Marawi/ Hebron | $37.88 \pm 3.46^{ef}$ | $1.64 \pm 0.26^{abc}$ |
| Shami/ Halhul | $36.86 \pm 7.27^f$ | $1.18 \pm 0.31^c$ |
| Shami/ Hebron | $46.66 \pm 1.25^{de}$ | $1.82 \pm 0.24^{ab}$ |
| p-value (ANOVA) | 0.00 | 0.03 |
| Bartlett's X2 | 9.69 | 8.67 |

Means separation following the ANOVA test was conducted using Tukey's HSD; values with a p-value $\leq 0.05$ are considered significant and have different letters.

The highest antioxidant activity was detected in the leaves of the Eshuhki cultivar from Hebron, followed by those of the Halawani cultivars from Al-Khadr and Hebron. In contrast, the lowest values were recorded for the leaves of the Baituni, Shami, and Marawi genotypes across all collection regions. These findings suggest that genotype is the primary factor influencing antioxidant capacity. For example, Halawani generally presented high antioxidant levels, whereas Baituni presented comparatively lower values. However, environmental conditions appear to play a modulatory role, as demonstrated by the significant variation in the antioxidant capacity among Halawani samples grown in different regions.

The results of the total phenolic content (TPC) revealed moderate yet statistically significant differences (p = 0.024) among the leaf samples. The highest TPC levels were detected in the leaves of Halawani (Halhul), Jandali (Al-Khadr), and Eshuhki (Hebron), whereas the lowest values were detected in the leaves of Shami (Halhul), Halawani (Hebron), and Baituni (Al-Khadr). Notably, the TPC values do not correlate with the antioxidant capacity values.

### 3.2 Metabolite profiling

Principal component analysis (PCA) clearly revealed that the collection region had no significant influence on metabolite concentrations (Figure 1 (A)). In contrast, a clear influence of the genotypes was evident, as the genotypes segregated based on their profiles of secondary metabolites (Figure 1 (B)).

**Figure 1.**
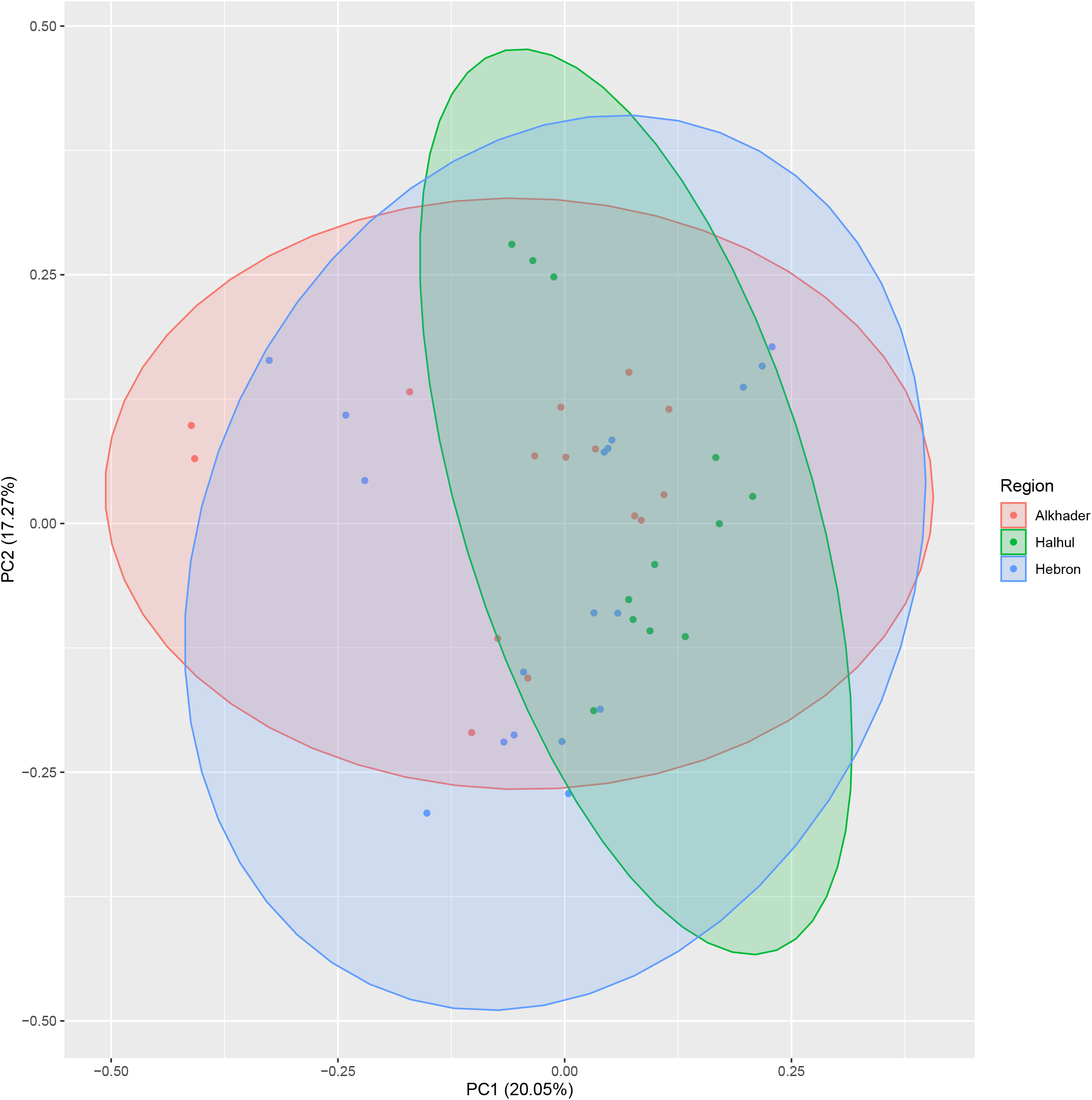
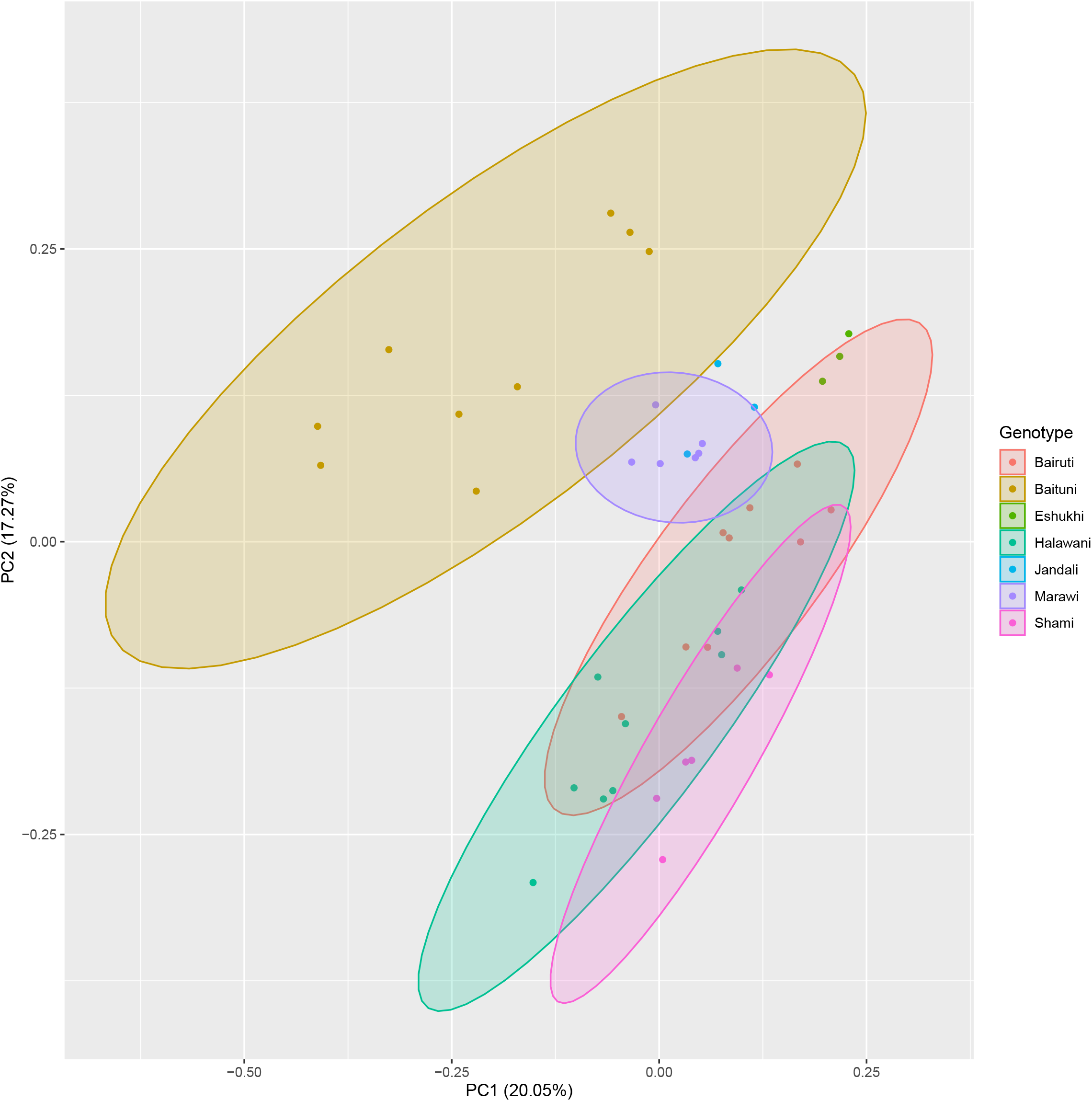
PCA of the effects of collection region (A) and genotype (B) on the secondary metabolites in grape leaves.

To understand the relationships between the original variables and principal components, a loading diagram was created to show how much each original variable contributed to each principal component (Figure 2).

**Figure 2.**
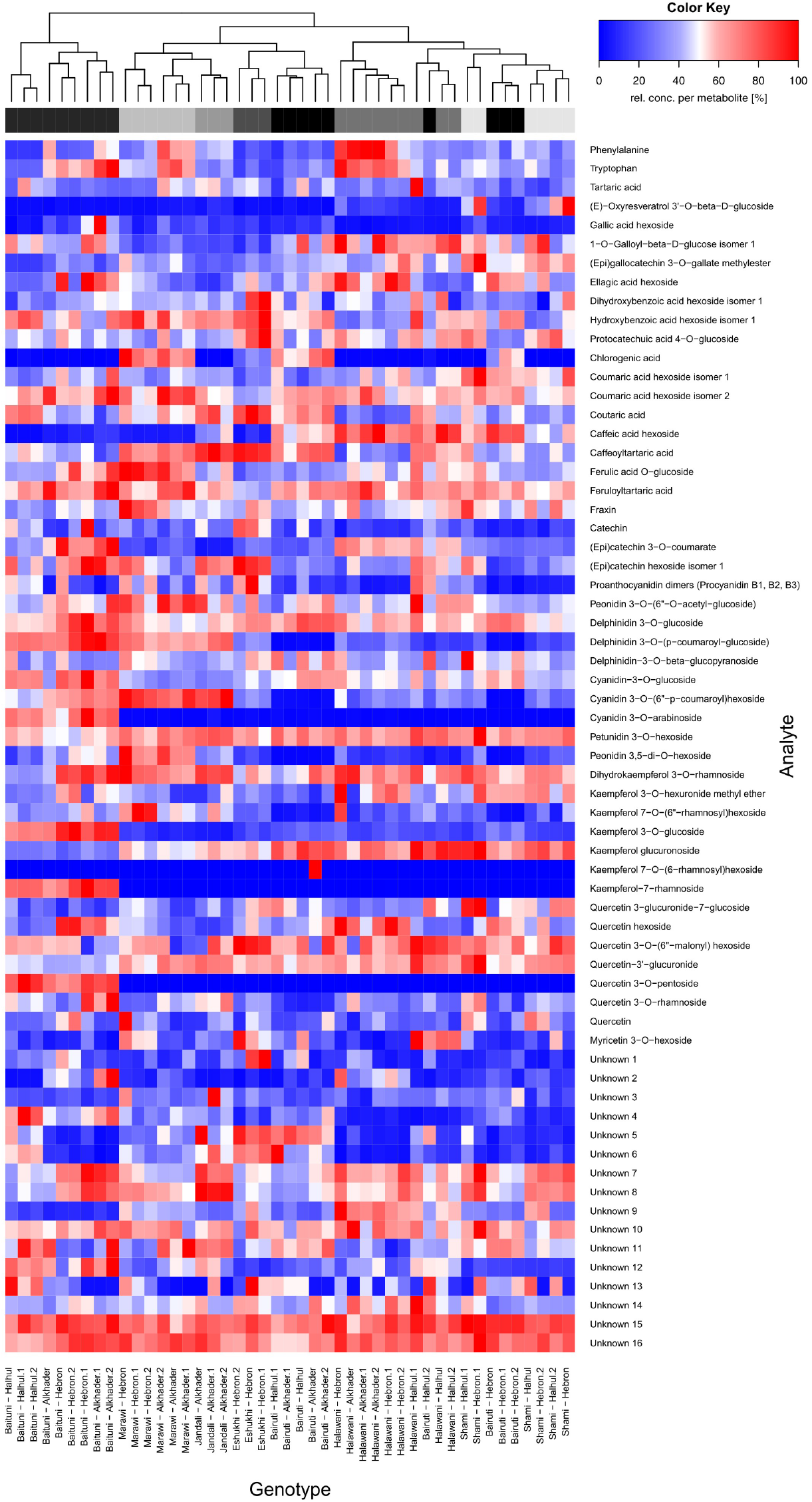
Hierarchical clustering and heatmap of secondary metabolites in grapevine leaves affected by genotype and collection region.

**Figure 3.**
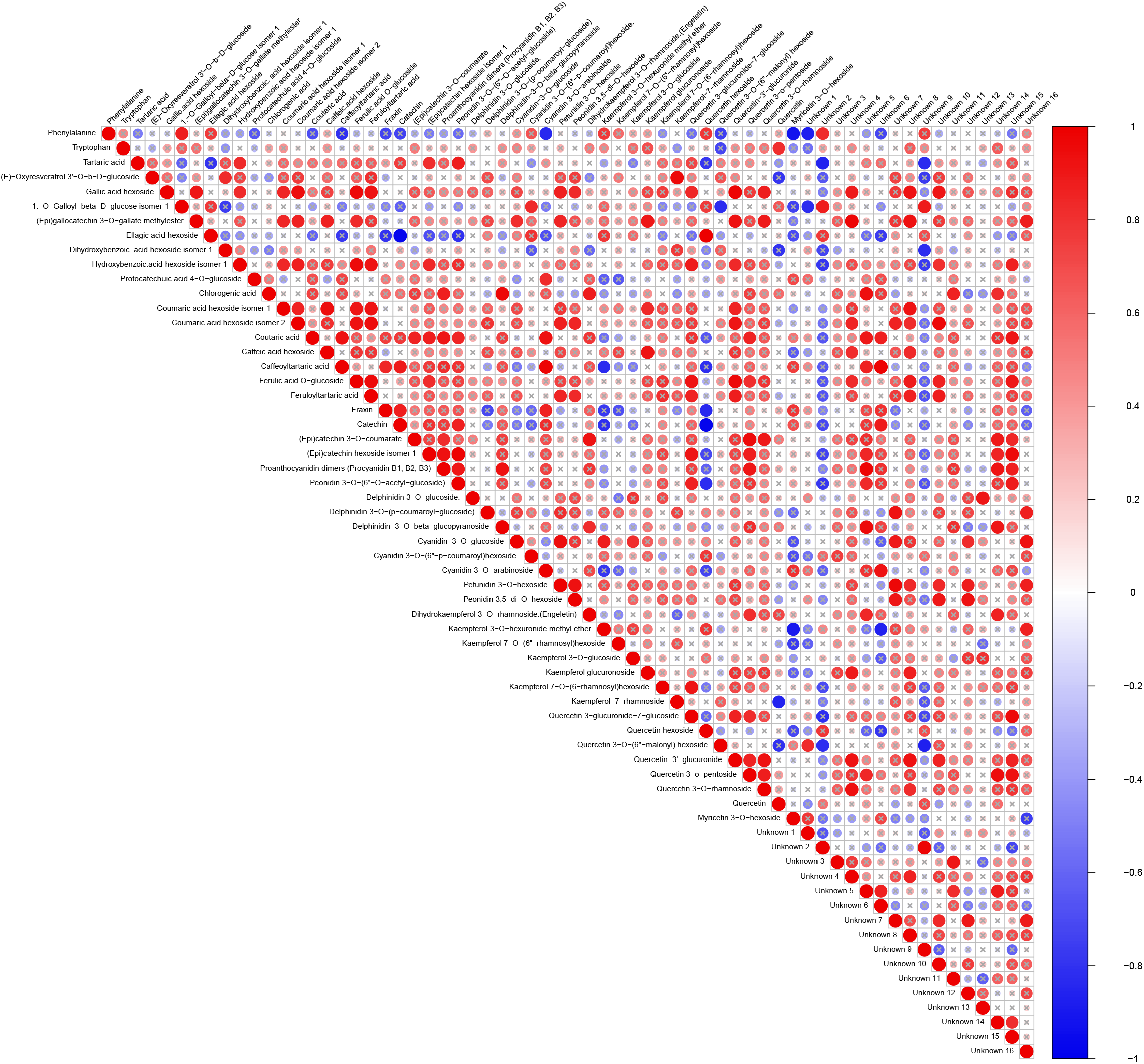
Pairwise correlation analysis of metabolite abundance in ‘Shami’ grapevine leaves.

**Figure 4.**
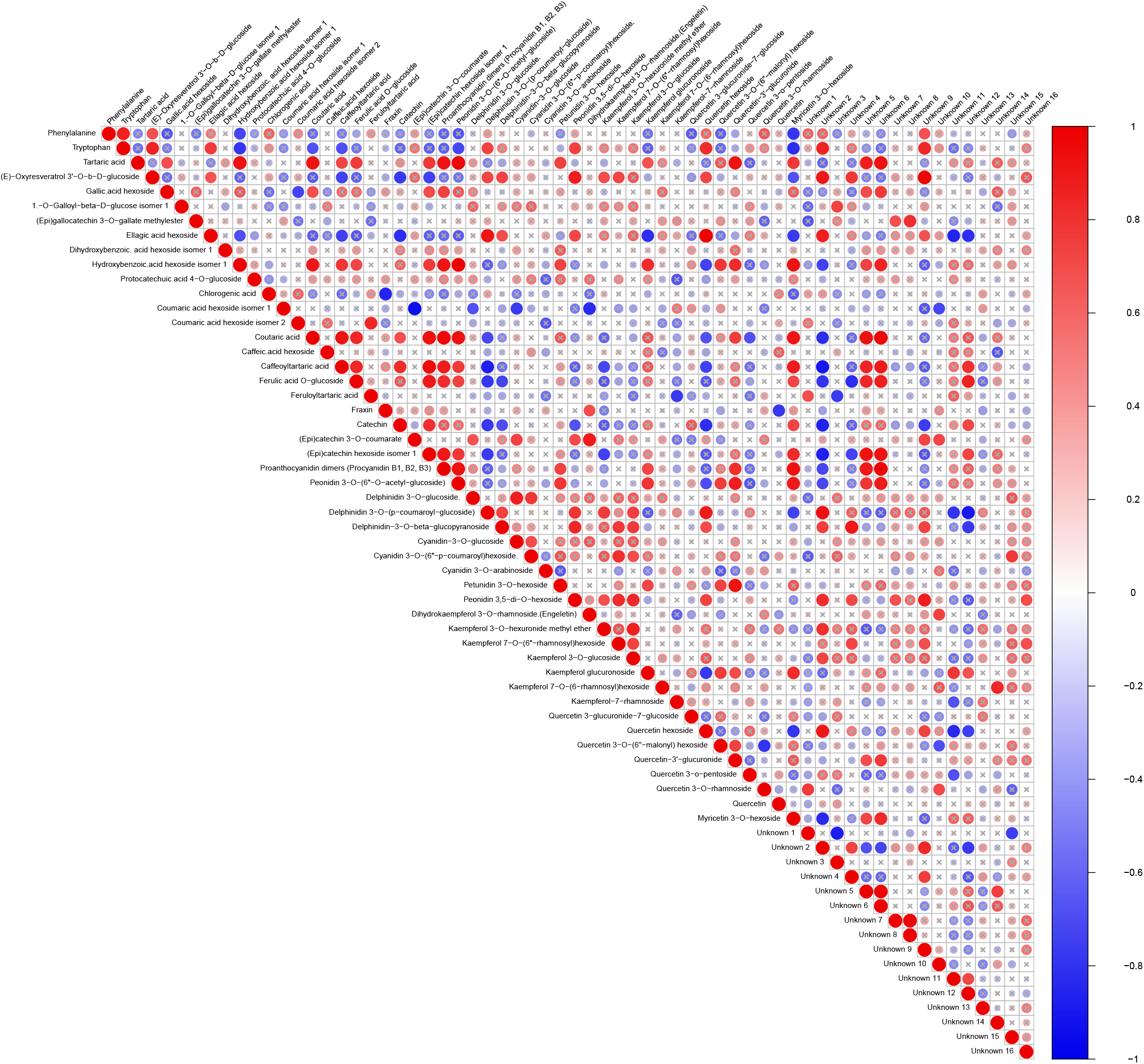
Pairwise correlation analysis of metabolite abundance in ‘Halawani’ grapevine leaves.

**Figure 5.**
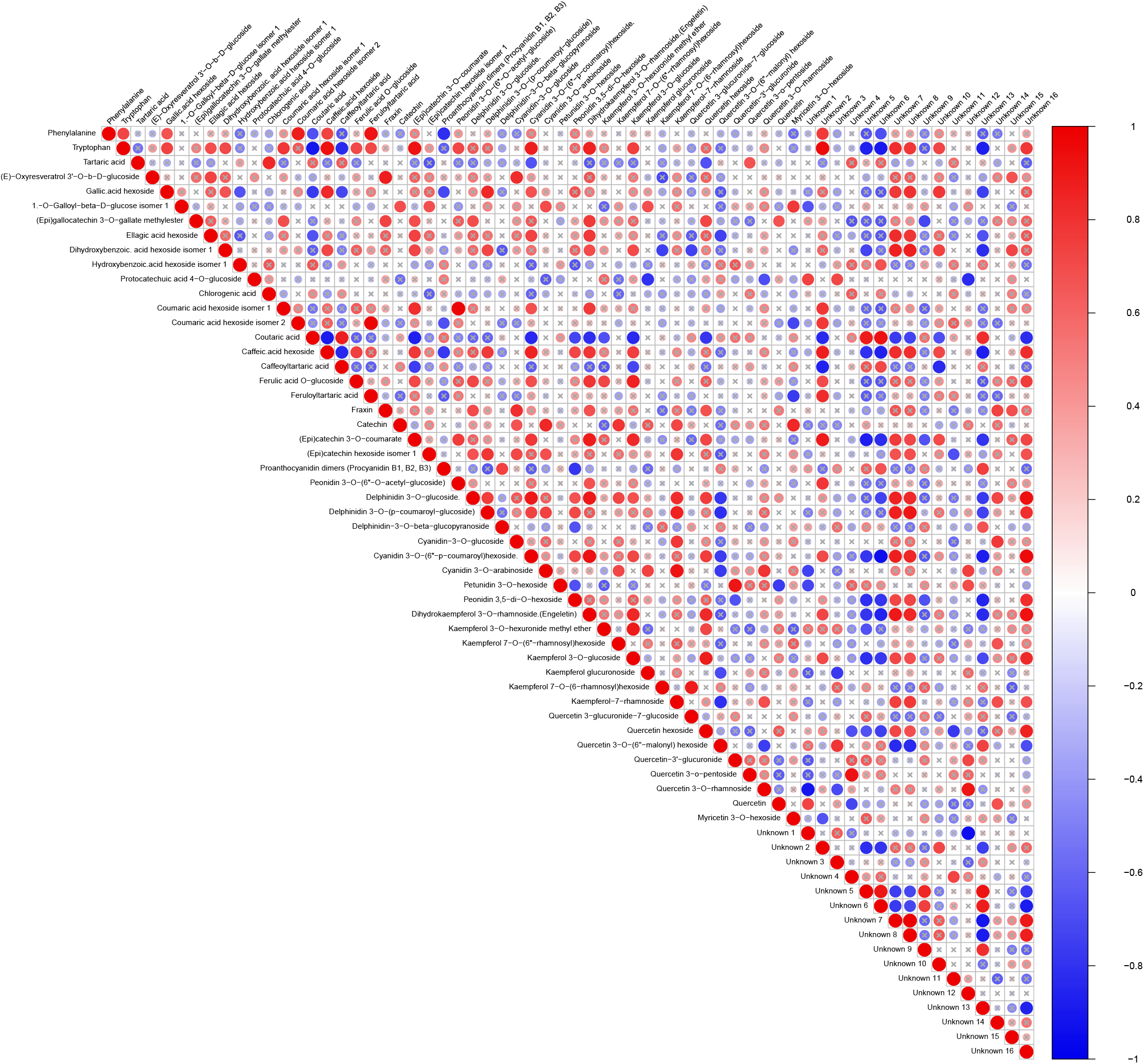
Pairwise correlation analysis of metabolite abundance in ‘Baituni’ grapevine leaves.

**Figure 6.**
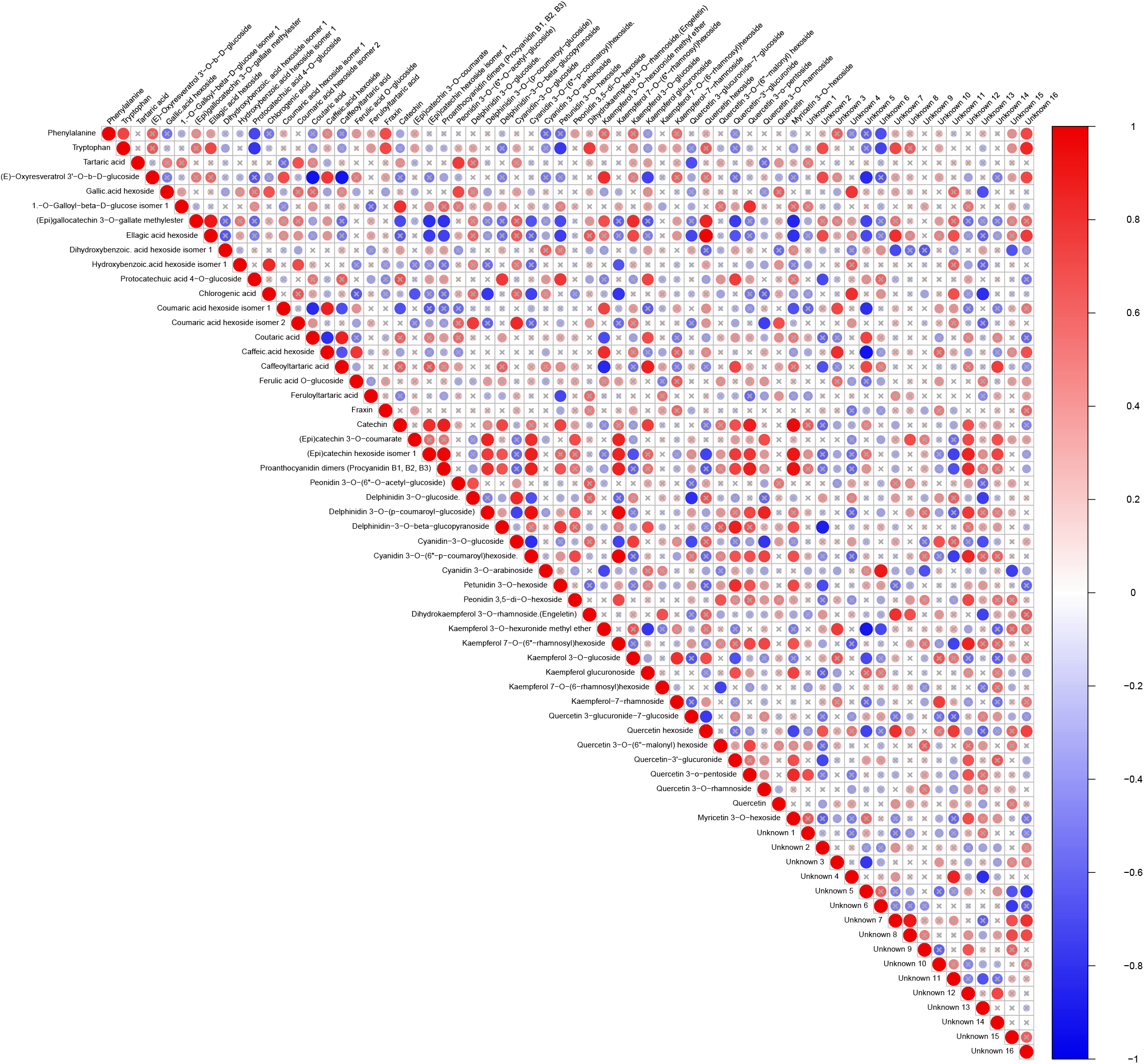
Pairwise correlation analysis of metabolite abundance in ‘Bairuti’ grapevine leaves.

**Figure 7.**
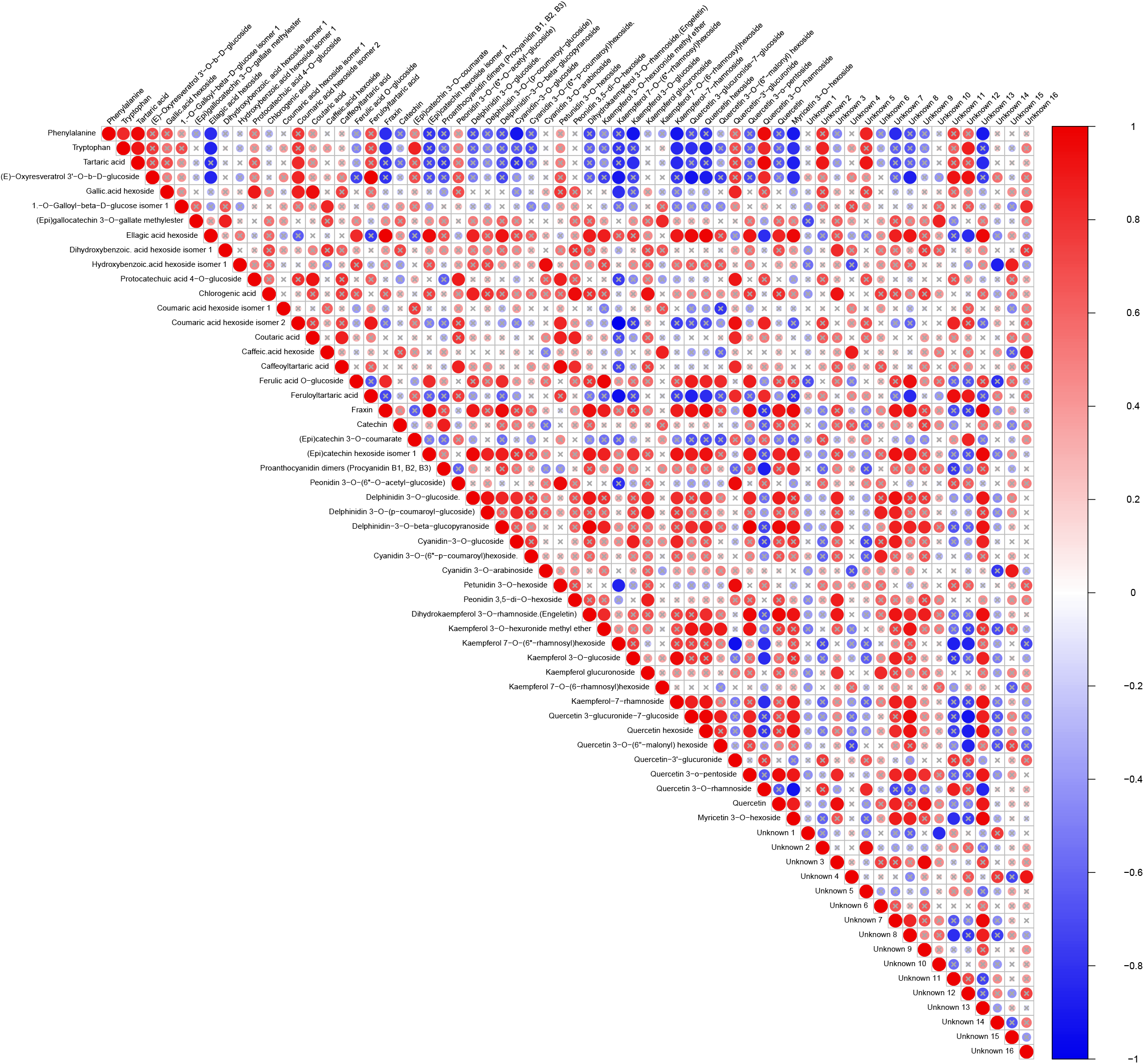
Pairwise correlation analysis of metabolite abundance in ‘Marawi’ grapevine leaves.

On the basis of their metabolic profiles, as well as pairwise correlation analysis of metabolite abundance, which was conducted for five of the seven assessed genotypes that were collected from more than one location, the grapevine genotypes covered in this study could be sorted into the following clusters:

#### Cluster 1 (Shami and Halawani)

Compared with other genotypes, the leaves of both genotypes presented fewer metabolites with elevated or moderate levels. Among these compounds are kaempferol glucuronoside and unknown 9 (m/z 482+). In contrast, a large set of metabolites were present at noticeably low levels. Among these compounds are quercetin hexoside, kaempferol-7-rhamnoside, quercetin, cyanidin 3-O-(6’’-p-coumaroyl)hexoside, delphinidin 3-O-(p-coumaroyl-glucoside), chlorogenic acid, coutaric acid, catechin, proanthocyanidin dimers, and unknown 5 (m/z 623-), unknown 6 (m/z 607-), unknown 7 (m/z 431-), and unknown 12 (m/z 507-). This set of metabolites can serve as a fingerprint for both genotypes. There are a number of metabolites that are specific to each genotype. For Shami grapes, gallic acid hexoside and (E)-oxyresveratrol 3’-O-b-D-glucoside are biosynthesized at high levels, whereas Halawani leaves contain high levels of caffeic acid hexoside, protocatechuic acid 4-O-glucoside, hydroxybenzoic acid hexoside isomer 1,1-O-galloyl-beta-D-glucose isomer 1, and unknown 2 (m/z 283-), unknown 3 (m/z 419-), and unknown 9 (m/z 482+).

#### Cluster 2 (Baituni grape)

Leaves of the Baituni grape give a unique profile. Although no single secondary metabolite can serve as a biomarker, a set of metabolites can easily distinguish this genotype. Among these are metabolites that are present at high to very high levels as well as metabolites with low to very low levels. The highly elevated metabolites included quercetin (3.15x), kaempferol-7-rhamnoside (3.37x), quercetin hexoside (1.70x), and quercetin 3-O-rhamnoside (1.35x). In addition, several unknowns can be included in this set (Unknown 10 (m/z 477-); Unknown 11 (m/z 407-); Unknown 12 (m/z 507-); and Unknown 13 (m/z 483-). The metabolites with the lowest levels included kaempferol glucuronoside, quercetin-3’-glucuronide, delphinidin-3-O-beta-glucopyranoside, caffoyltartaric acid, caffeic acid hexoside, ellagic acid hexoside, catechin, (E)-oxyresveratrol 3’-O-b-D-glucoside, proanthocyanidin dimers, unknown 5 (m/z 623-), and unknown 9 (m/z 482+). Interestingly, all flavonol glycosides either maintain their levels or increase drastically, except for two flavonol glycosides: kaempferol glucuronoside and quercetin-3’-glucuronide.

#### Cluster 3 (Eshukhi, Bairuti)

Leaves of both genotypes presented elevated levels of caffeoyltartaric acid, coutaric acid, ferulic acid O-glucoside, and proanthocyanidin dimers; unknown 5 (m/z 623-), unknown 6 (m/z 607-), and unknown 14 (m/z 537+). On the other hand, the levels of quercetin hexoside, kaempferol-7-rhamnoside, quercetin, dihydrokaempferol 3-O-rhamnoside, cyanidin 3-O-(6’’-p-coumaroyl)hexoside, and peonidin 3-O-(6’’-O-acetyl-glucoside) are lower than those in the other genotypes. The above-listed metabolites can serve as a fingerprint for both genotypes. In addition, a complementary set of metabolites for each genotype was generated. The complementary set for Eshukhi grapes included the following metabolites that exist at noticeably high levels: tartaric acid, coumaric acid hexoside isomer 1, and catechin. The set for Bairuti includes delphinidin-3-O-beta-glucopyranoside, chlorogenic acid, caffeic acid hexoside, ellagic acid hexoside, and unknown 13 (m/z 483-).

#### Cluster 4 (Jandali, Marawi)

Leaves of both genotypes contain high levels of cyanidin 3-O-(6’’-p-coumaroyl)hexoside, delphinidin 3-O-(p-coumaroyl-glucoside), quercetin hexoside, peonidin 3-O-(6’’-O-acetyl-glucoside), dihydroxybenzoic acid hexoside isomer 1, proanthocyanidin dimers, unknown 2 (m/z 283-), and unknown 3 (m/z 419-). In contrast, they contain low levels of kaempferol-7-rhamnoside and quercetin. Complementary to these metabolites, which constitute a set that dictates the fingerprint for both genotypes, for each genotype, there is a set of metabolites that add more specificity to the finger for each; these metabolites exist at elevated levels. For Marawi grapes, the complementary set included chlorogenic acid, protocatechuic acid 4-O-glucoside, ellagic acid hexoside, hydroxybenzoic acid hexoside isomer 1, and unknown 9 (m/z 482+). The complementary set for Jandali grapes included caffeoyltartaric acid, catechin, caffeic acid hexoside, unknown 11 (m/z 407-), and unknown 12 (m/z 507). Notably, both Jandali and Marawi. In contrast to Eshukhi, Bairuti, Halawani, and Shami, but similar to Baituni, it contains high levels of cyanidin 3-O-(6’’-p-coumaroyl)hexoside and delphinidin 3-O-(p-coumaroyl-glucoside).

## 4. Discussion

Staffed grapevine leaves are a staple dish across the Mediterranean region (Ali-Shtayeh et al., 2018). Notably, this cuisine is highly appreciated not only for its taste but also for its nutritional value due to its high polyphenol content (Yang and Xiao, 2013). The significance of the metabolic profiling of the leaves of grapes performed in this study is far beyond assessing the nutritional quality of grape leaves, as it can be a very valuable tool for chemotaxonomic purposes and future breeding programs.

Our results revealed that the antioxidant capacity and total phenolic content of leaves significantly differ among the genotype/location combinations. The leaves of Eshuhki grape collected from Hebron outperformed those of the other genotypes in terms of both parameters. The reasons behind this are not clear, although it is possible that genetic factors play a role. In addition, no specific location shows a pattern on its own, with either lower or higher values. Therefore, we believe that the microclimate within each vineyard is the decisive factor. In this sense, we do not believe that acute drought plays a role, since the picking time in the spring season ensures that soil moisture is still adequate for grapevines, which are cultivated without supplementary irrigation. However, the same grapevines in these regions experience significant drought in summer during the berry-ripening period. Accordingly, we believe that adaptation to harsh environments in summer might have a profound effect on the metabolic profiles of leaves, even before the onset of any stress. It is logical to believe that plants initiate adaptive responses to anticipated abiotic stress or stresses early in the growing season, and the localization of such evolutionary adaptive responses will logically be in roots and leaves. In this respect, Ding et al. (2020) reported that plants detect early-season signals, such as photoperiodic signals and temperature trends, that predict future stresses.

Considering that differences between genotypes are profound and significant, in contrast to location, our focus here will be on these genotypes. In this respect, the current major challenges in viticulture are drought (Keller, 2023) and powdery mildew infection (Sosa-Zuniga et al., 2022). Accordingly, elucidating any possibility that helps in counteracting the negative impacts of drought and powdery mildew disease is highly important.

We begin with drought-tolerant grapes, and our hypothesis is that some old indigenous grape genotypes (e.g., Eshukhi) are more drought-tolerant than are newly introduced grape genotypes (e.g., Halawani). This hypothesis is based fully on interviews with local farmers and agricultural extension officers. To identify metabolites that might contribute to drought tolerance in Eshukhi compared with Halawani, we compared the metabolic profiles across both genotypes. The most noticeable difference is the profound accumulation of specific protective metabolites that are crucial for abiotic stress tolerance (Sharma et al. (2019)). The main group of metabolites that are found in the drought-tolerant genotype at much higher levels are hydroxycinnamic acids. These acids and their derivatives were found to increase under stressful conditions as a means to withstand environmental stresses (Khawula et al., 2023). In addition, a study with barley confirmed the accumulation of hydroxycinnamic acid derivatives under drought conditions (Piasecka et al., 2017). In this sense, caffeoyltartaric acid is a caffeic acid derivative that is a powerful antioxidant that is biosynthesized massively under drought stress to avoid cellular damage (Nakabayashi et al., 2014). In addition to hydroxycinnamic acid derivatives, catechin is highly elevated in drought-tolerant genotypes. A previous study revealed that the total amount of catechin increased under drought stress conditions and that the accumulation of catechin under such conditions included the polymerization of proanthocyanidins, which scavenge excess reactive oxygen species (Lv et al., 2021). Furthermore, Xu et al. (2025) reported that many biosynthetic genes associated with catechins are induced under stressful conditions, which results in increased catechin levels under abiotic stress.

For the second major challenge, powdery mildew, the comparison is between Marawi and Jandali, which are believed to be more tolerant to powdery mildew, and Bairuti, which is very sensitive to powdery mildew. The profound differences in the metabolic profile of the abovementioned genotype are obvious in the levels of many metabolites, including quercetin hexoside, catechin, caffeoyltartaric acid, petunidin 3-O-hexoside, dihydroxybenzoic acid hexoside isomer 1, fraxin, dihydrokaempferol 3-O-rhamnoside, proanthocyanidin dimers, and the unknown 1 (m/z 427-). These metabolites might contribute to increased resistance to powdery mildew (Dixon, 2001). A study of healthy and diseased grape leaves (infected with Erysiphe necator) revealed that the levels of anthocyanidins, hydroxycinnamic acids (mainly feruloyl derivatives), and epigallocatechin gallate were greater in infected leaves than in infected leaves; coumaric acid, feruloyl tartaric acid, peonidin hexoside, and coumaric derivatives of peonidin and cyanidin were more abundant (Hernández et al., 2024).

The importance of secondary metabolites in grape leaves extends beyond tolerance to drought stress and powdery mildew infection: these metabolites contribute to nutritional quality and taste. On this basis, we address these two genotypes separately. The **Shami** grape is one of the appreciated grapes in PS and produces highly pigmented berries. The large number of anthocyanin glycosides and flavonol glycosides present at normal levels may indicate the strong pigmentation potential of the leaves, although they are masked by their green color. Interestingly, high levels of (E)-oxyresveratrol 3’-O-b-D-glucoside and gallic acid hexoside affect health (Navarro et al., 2018; Galindo et al., 2011; Šuković et al., 2020).

**Marawi** is another beautiful genotype that is believed to be one of the ancient grape genotypes in historical PS and was recently at the center of a political dispute (Monterescu and Handel, 2019). The leaves of Marawi grape presented notably high levels of chlorogenic acid, which was among the highest observed across genotypes. From a health perspective, this is significant because of the well-known antioxidant and anti-inflammatory properties of this acid (Yun et al., 2012).

## 5. Conclusion

The present study revealed the diversity of grape genotypes at the metabolomic level. This is a first step toward a comprehensive research effort to collect, investigate, and assess the complexity of grape culture in PS, which is one of the regions with ancient viticulture. Future efforts will employ a holistic approach that involves, among other methods, genomic, transcriptomic, and metabolomic analyses. The final goal is to preserve the current genotypes and develop grape varieties that can cope with the adverse impacts of climate change and improve the health of people in marginalized regions.

## Supporting information

Cover letter

Ethical statement

## Acknowledgment

The financial support of the International Development Research Centre (IDRC) – Canada is highly appreciated.

