## Supplementary material for "Profiling of Secondary Metabolites of Leaves of Indigenous and Introduced Grapes Collected from Hebron and Bethlehem Regions in the West Bank-Palestine": Cover letter

**Date:** 04-02-2026

Dear Editor,

I am submitting, as the corresponding author, the above mentioned research article, which is part of a research project that addressed consumption of fruits and vegetables in Palestine funded by IDRC – Canada.

The significance of the submitted study lies on the importance on grapevine leaves as a stable foodstuff in the Mediterranean, particularly in the marginalized rural areas. In addition, we strongly believe that assessing the metabolic profiles of indigenous grape genotypes is of great value to distribute such old genotypes, as well as to develop new varieties that withstand the harsh abiotic stresses, in particular drought. Furthermore, we are highly interested in understanding the mechanisms of the tolerance of few old grape genotypes to powdery mildew infection, which is a very devastating fungal disease in Palestine and other Mediterranean countries.

We are planning, in cooperation with international partners, to proceed working with old indigenous grape genotypes, which will adopt a more holistic approach. This approach will include genomics, transcriptomic, and further metabolomics assessments.

Please be aware also of the following two documents related to the submitted article that are uploaded as supplementary documents:

Supplementary file 1: the IRB approval, which was issued before we started the study.

Supplementary file 2: the description of the statistical analysis.

Sincerely yours

**Jamil Harb**

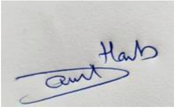
