## Supplementary material for "Profiling of Secondary Metabolites of Leaves of Indigenous and Introduced Grapes Collected from Hebron and Bethlehem Regions in the West Bank-Palestine": Ethical statement

معهـــد الصحـــة العامـــة والمجتمعيـــة
Institute of Community and Public Health

جامعة بيرزيت
BIRZEIT UNIVERSITY

**Institute of Community and Public Health – Birzeit University**
**Ethics Review Committee Decision**

**Date**: January 31, 2019
**Applicant's name**: Dr. Abdullatif Husseini
**Institution:** Institute of Community and Public Health

| | |
|---|---|
| **Reference No:** | **2019 (1 – 3)** |
| **Project Title:** | Improving food policies and enabling healthier diets for preventing non-communicable diseases in the West Bank |
| **Names of contributing researchers, other than the Principal Investigator/ Applicant:** | Abdullatif Husseini ICPH-BZU<br>Jamil Harb Department Of Biology And Biochemistry - BZU<br>Rita Giacaman ICPH-BZU<br>Rawan Kafri ICPH-BZU<br>Dalia Ghosheh ICPH-BZU |

Thank you for submitting your application for the ethics review of your research proposal. Your application was examined carefully, and discussed by the Ethics Review Committee during a meeting which took place on January 31, 2019. The following documents were reviewed:

1. Ethics Review Application Letter/Form
2. Consent Form
3. Project proposal and related documents, as revised based on the comments of the Committee.
4. Other forms and permissions (*as necessary*)

**The ICPH-BZU Research Ethics Review Committee has approved your research proposal.**
Approval is given for three years. Projects, which have not commenced within two years of original approval, must be re-submitted to the Ethics Review Committee. You must inform the Committee when the research has been completed. If you are unable to complete your research within the five year validation period, you will be required to write to the Ethics Review Committee to request an extension. You may also need to re-apply for approval by the Committee.
Any serious adverse events or significant changes which occur in connection with this study and/or which may alter its ethical considerations must be reported immediately to the Ethics Review Committee. On such an occasion, an "Amendment Form" must be submitted to the Committee for re-assessment.

Thank you,
Ethics Review Committee Coordinator
**Maysaa Nemer, PhD**

جامعة بيرزيت Institute of Community & Public Health

معهـــد الصحـــة العامـــة والمجتمعيـــة
Institute of Community and Public Health

جامعة بيرزيت
BIRZEIT UNIVERSITY

**Institute of Community and Public Health – Birzeit University**
**Ethics Review Committee (ERC)**

**Part I: To be completed by the applicant:**

**Date of application**: January 10, 2019
**Applicant's name**: Dr. Abdullatif Husseini
**Institution:** Institute of Community and Public Health

| | |
|---|---|
| **Reference No:** | **2019 (1 – 3)** |
| **Project Title:** | Improving food policies and enabling healthier diets for preventing non-communicable diseases in the West Bank |
| **Names of contributing researchers, other than the Principal Investigator/ Applicant:** | Abdullatif Husseini ICPH-BZU<br>Jamil Harb Department Of Biology And Biochemistry - BZU<br>Rita Giacaman ICPH-BZU<br>Rawan Kafri ICPH-BZU<br>Dalia Ghosheh ICPH-BZU |

Comments: (if any)

**Part II: To be completed by the (ERC)**

***Decision:**
**The ICPH-BZU Research Ethics Review Committee approves this study.**

*** Detailed information will be provided in a separate form.**

***Decision date: January 31, 2019***

**Ethics Review Committee members, qualifications and signatures:**

| Name | Specialty | Qualifications | Signature |
|---|---|---|---|
| Niveen Aburmeileh | Statistical Epidemiology | PhD | Niveen AbuRmeileh |
| Suzan Mitwalli | Public Health | MPH | [signature] |
| Shiraz Nasr | Spatial analysis | MSA | [signature] |

جامعة بيرزيت Institute of Community & Public Health
